# A ROLE FOR CORTICAL DOPAMINE IN THE PARADOXICAL CALMING EFFECTS OF PSYCHOSTIMULANTS

**DOI:** 10.1101/2021.10.01.462763

**Authors:** Sharonda S. Harris, Sara M. Green, Mayank Kumar, Nikhil M. Urs

**Affiliations:** Department of Pharmacology and Therapeutics, University of Florida, Gainesville FL 32610

## Abstract

Attention deficit hyperactivity disorder (ADHD) affects young children and manifests symptoms such as hyperactivity, impulsivity and cognitive disabilities. Psychostimulants, which are the primary treatment for ADHD, target monoamine transporters and have a paradoxical calming effect, but their mechanism of action, is unclear. Studies using the dopamine (DA) transporter (DAT) knockout mice, which have elevated striatal DA levels and are considered an animal model of ADHD, have suggested that the paradoxical calming effect of psychostimulants might be through the actions on serotonin neurotransmission. On the other hand, newer non-stimulant class of drugs such as atomoxetine and Intuniv suggest that targeting the norepinephrine (NE) system in the PFC might explain this paradoxical calming effect. We sought to decipher the mechanism of this paradoxical effect of psychostimulants through an integrated approach using *ex vivo* monoamine efflux experiments, monoamine transporter knockout mice, drug infusions and behavior. Our *ex vivo* efflux experiments reveal that NE transporter (NET) blocker desipramine elevates both norepinephrine and dopamine but not serotonin levels, in PFC tissue slices from wild-type and DAT-KO but not NET KO mice. However, serotonin (5-HT) transporter (SERT) inhibitor fluoxetine elevates only serotonin in all three genotypes. Systemic administration of both desipramine and fluoxetine but local PFC infusion of only desipramine and not fluoxetine inhibits hyperactivity in the DAT-KO mice. In contrast, pharmacological norepinephrine depletion but dopamine elevation using Nepicastat also inhibits hyperactivity in DATKO mice. Together, these data suggest that elevation of PFC dopamine and not norepinephrine or serotonin as a convergent mechanism for the paradoxical psychostimulant effects observe in ADHD therapy.

## INTRODUCTION

Attention deficit hyperactivity disorder (ADHD) affects at least 5-10% of children, with increasing diagnoses every year. The primary line of treatment for ADHD are the psychostimulant class of drugs including amphetamine (Aderall) and methylphenidate (Ritalin). Several theories have been postulated on the paradoxical calming effect of psychostimulants on ADHD patients but even after decades of research, their exact molecular and anatomical mechanism of action is not clear. Studies regarding the role of different monoamine transporters in the mechanism of action of psychostimulants have been inconclusive. Studies using hyperactive dopamine transporter (DAT) knockout (DAT-KO) mice suggest a serotonergic mechanism since serotonin transporter (SERT) inhibitor fluoxetine or serotonin precursor 5-HTP inhibit hyperactivity (Gainetdinov et al., 1999; Giros et al., 1996). However other studies using microdialysis and fluoxetine suggest a role for cortical norepinephrine and dopamine (Bymaster et al., 2002; Tanda et al., 1994). Additionally, newer non-stimulant forms of ADHD medications (atomoxetine, Intuniv) disrupt norepinephrine (NE) transmission, thus providing evidence for a role for the NE transporter (NET) in this paradoxical calming effect. Interestingly, NET knockout (NET-KO) mice have reduced locomotor activity and lower striatal DA release (Xu et al., 2000), again suggesting a crucial role for NET in this calming effect. However, NET inhibition not only elevates norepinephrine levels but also dopamine levels in the PFC (Moron et al., 2002; Tanda et al., 1994; Xu et al., 2000). Therefore, it is still not clear whether PFC norepinephrine, dopamine or serotonin are necessary for this paradoxical effect.

We hypothesized that PFC dopamine is the common mechanism that drives this paradoxical effect. Previous microdialysis studies have established an inverse relationship between DA levels in the PFC and striatal DA levels, and locomotor activity (Ventura et al., 2004a; Ventura et al., 2004b). DA depletion by pharmacological lesioning in the PFC has been shown to increase subcortical dopamine and amphetamine-induced locomotor activity compared to non-lesioned animals (Pycock et al., 1980; Sokolowski and Salamone, 1994; Ventura et al., 2004a). Additionally, we have previously shown that D2 receptor deletion in the PFC increases PCP-induced hyperactivity (Urs et al., 2016).

The present study investigated the effects of pharmacological agents that increase DA levels in the PFC on locomotor activity. Increasing DA levels systemically or locally in the PFC either by NET inhibition or dopamine beta hydroxylase (DBH) inhibition led to a reduction in DATKO hyperactivity. The data from the current study provide a mechanistic insight into the regulation of monoamines and the paradoxical effects of psychostimulants.

## RESULTS

NET-KO mice have reduced locomotor activity (Xu et al., 2000), whereas SERT inhibition reduces hyperactivity in the DAT-KO mice (Gainetdinov et al., 1999). We first sought to test the effects of NET or SERT inhibition in the DAT-KO mice. Systemic administration of the NET inhibitor desipramine (25 mg/kg, i.p.) or fluoxetine (20 mg/kg, s.c.) significantly decreased hyperactivity in DATKO mice, compared to vehicle (saline) injected controls (**Figure 1**). Our findings are in contrast to the original study (Gainetdinov et al., 1999), which shows that NET inhibition using Nisoxetine does not affect hyperactivity in the DAT-KO mice.

**Figure 1.**
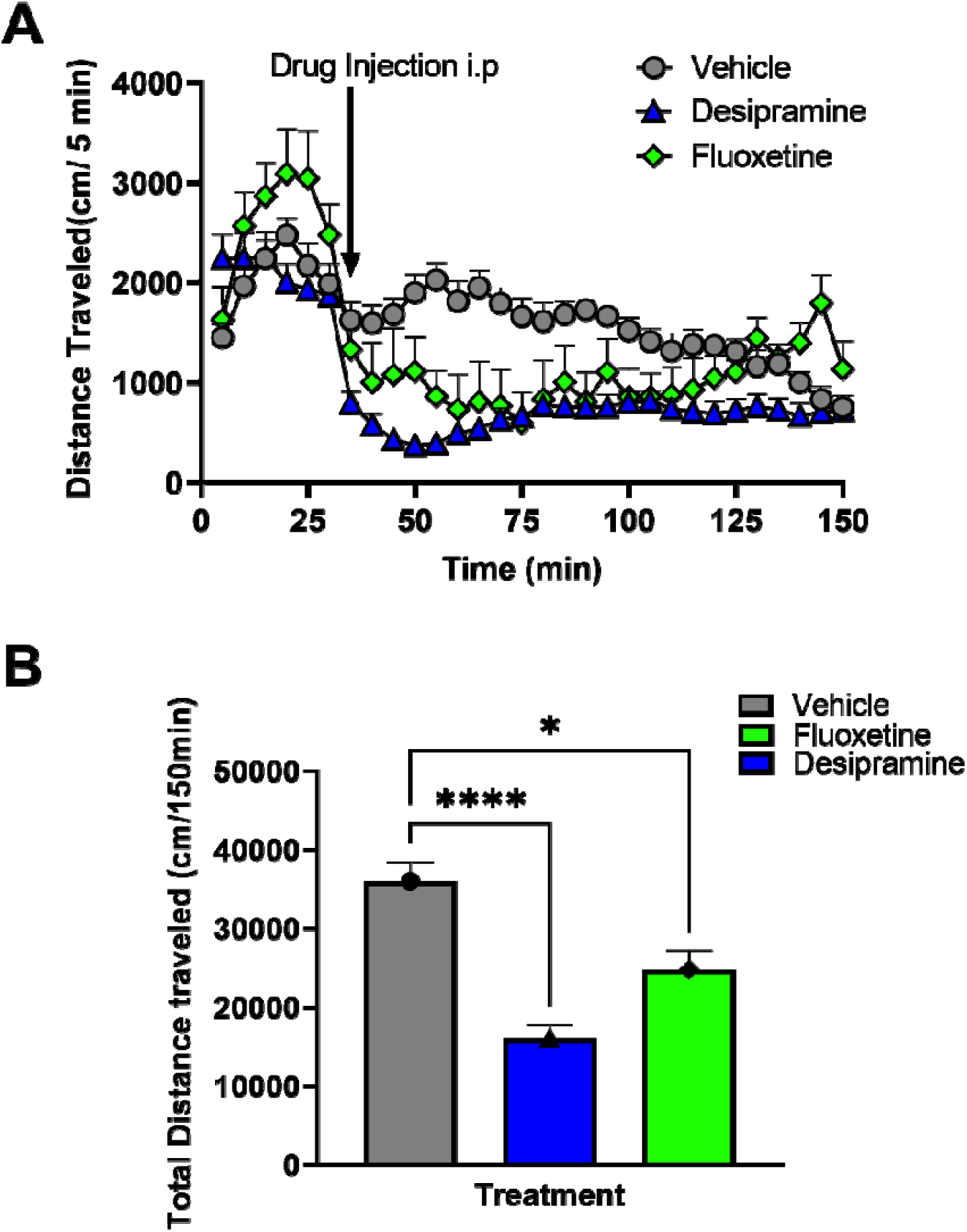
Effect of monoamine transporter drugs on DAT-KO hyperactivity. Hyperactive DAT-KO mice were systemically injected with vehicle (saline) or transporter blockers Desipramine (NET) or fluoxetine (SERT) and **A**. after a 30min baseline locomotor activity (cm/5min) was recorded for an additional 120min, or **B**. total distanc traveled (cm/150min) calculated. N=6-8 mice per group. *P<0.05, ****P<0.0001 using Two-Way ANOVA compared t Vehicle-treated controls.

The PFC is thought to be the site of action of psychostimulants and other therapeutics in ADHD therapy (Arnsten and Dudley, 2005; Berridge et al., 2006), since they improve motivational and cognitive functions. We asked the question if desipramine and fluoxetine alter monoamine levels in the PFC. To test this question, we used a previously established ex vivo efflux assay (Johnson et al., 2005; Pino et al., 2021), that allows measuring monoamines released from tissue slices submerged in a well with oxygenated buffer. Potassium chloride (KCl) application to tissue slices induces release of monoamines that can be measured by HPLC. The KCl-mediated monoamine release can be further enhanced by application of monoamine transporter drugs such as amphetamine (see Methods section for details). We first tested amphetamine-induced efflux in PFC tissue slices from WT, DAT, and NET KO mice. Amphetamine (Amph) significantly elevated only DA levels in DAT-KO mice but not WT or NET-KO mice (**Figure 2**). NE and 5-HT levels were unchanged in the WT and DAT-KO mice. However, in the NET KO mice there was a trend towards reduced NE and 5-HT levels upon Amph treatment. Amphetamine is non-specific for monoamine transporters and elevates monoamine levels by inducing reverse transport. We next tested specific transporter blockers desipramine, GBR12909, and fluoxetine that inhibit NET, DAT and SERT, respectively. In WT mice, NET inhibition with desipramine (10 uM) increased KCl-stimulated NE and DA compared to KCl alone (**Figure 3Ai, DA; 3Aii NE**). As expected, SERT inhibition with Fluoxetine (10 uM) increased KCl-stimulated 5-HT in WT mice, but did not alter NE or DA levels (**Figure 3Aiii, 5-HT**). Our findings are in contrast to previous studies using microdialysis that show that fluoxetine elevates NE and DA in the PFC (Bymaster et al., 2002; Tanda et al., 1994). GBR12909 had no effect in PFC tissue slices consistent with evidence that DAT expression is very low in the PFC compared to striatum and reuptake of dopamine is primarily mediated by the NET (Ciliax et al., 1995; Moron et al., 2002; Sesack et al., 1998). In DAT-KO mice, NET inhibition led to a significant increase in KCl-stimulated NE, DA, but not 5-HT levels in DATKO mice compared to KCl treatment alone (**Figure 3Bi, DA; 3Bii, NE**). The increase in DA levels in DAT-KO mice is much higher than that of WT mice. Similar to WT mice, in DAT-KO mice SERT inhibition with fluoxetine led to a significant increase in KCl-stimulated 5-HT levels but not NE or DA levels compared to KCl treatment alone. Lastly, in NETKO mice, NET or DAT inhibition had no effect on KCl-stimulated NE, DA, or 5-HT levels (**Figure 3Ci-iii**), whereas SERT inhibition increased only KCl-stimulated 5-HT but not NE or DA (**Figure 3Ci-iii, 5-HT**). The changes in monoamine release in DAT-KO mice are not due to homeostatic changes in PFC tissue monoamine levels, as shown by HPLC (**Figure S1 A, B**), but are consistent with lack of release in NET KO mice since tissue levels of NE are significantly lower (**Figure S1 C, D)**. Since our results show that systemic NET inhibition reduces hyperactivity in DAT-KO mice and that NET inhibitors elevate NE and DA levels in the PFC of DAT-KO mice, we next wanted to test the effect of PFC NET inhibition in DAT-KO mice. Bilateral infusion of desipramine (4ug/0.5 ul/side) significantly reduced hyperactivity of DAT-KO mice (**Figure 4A, B**). Consistent with our efflux data, infusion of fluoxetine did not significantly affect hyperactivity of DATKO mice (**Figure 4A, B**). Additionally, bilateral infusion of amphetamine (Amph, 100uM/0.5ul/side) also significantly reduced hyperactivity in DAT-KO mice (**Figure 4B**).

**Figure 2.**
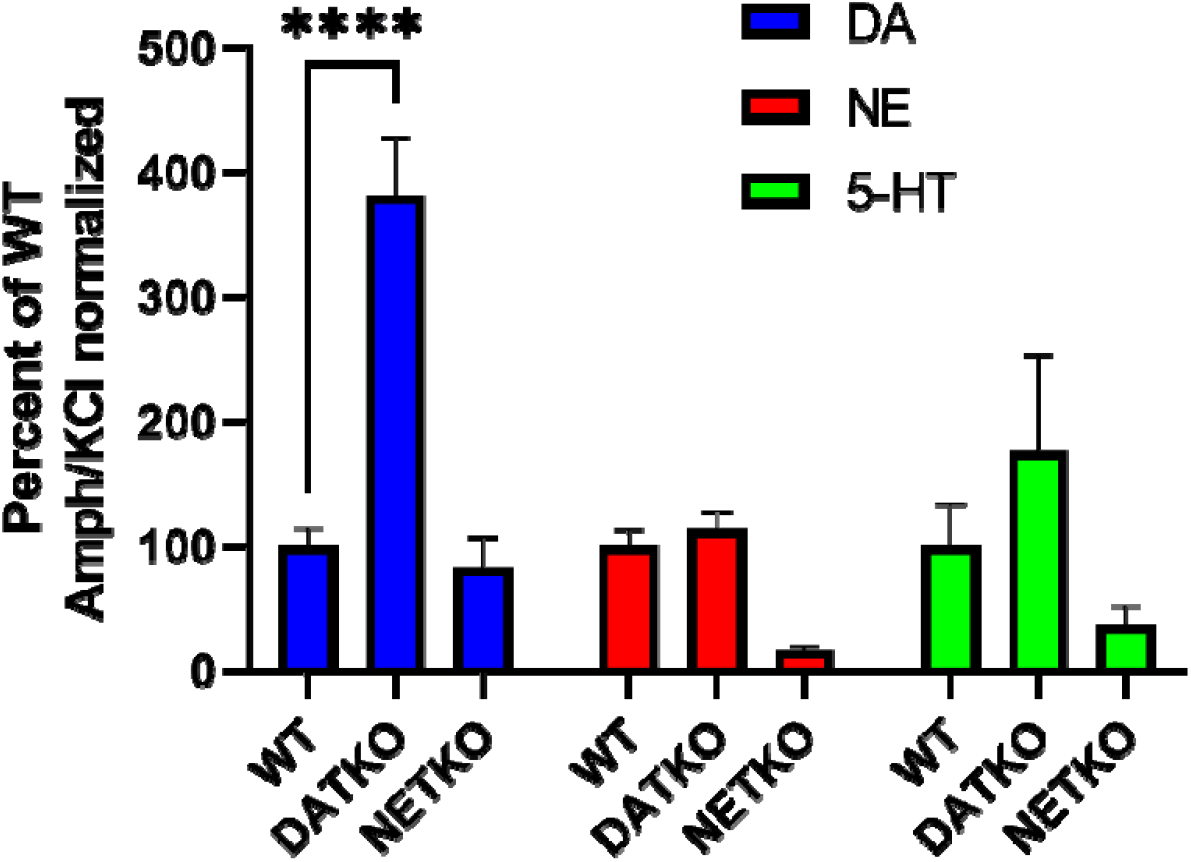
Effect of Amphetamine on monoamine efflux in PFC tissue. PFC tissue slices from WT, DAT-KO and DAT-KO mice were treated with amphetamine (AMPH, 10uM) and monoamines released into the incubation KH buffer were analyzed by HPLC. ****P<0.0001 using Two-Way ANOVA, compared to WT mice.

**Figure 3.**
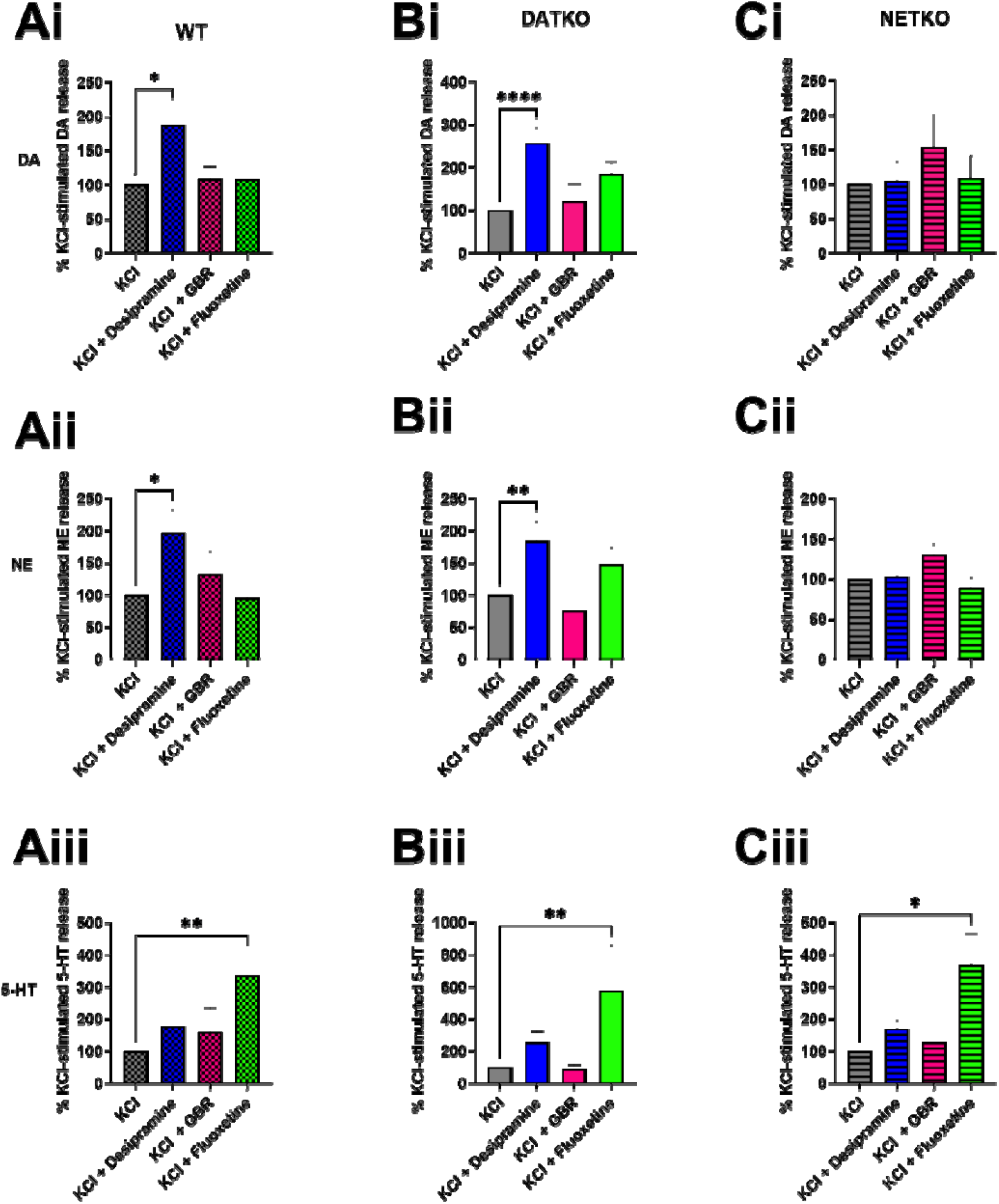
Effect of monoamine transporter blockers on monoamine efflux in PFC tissue. PFC tissue slices from WT, DAT-KO and NET-KO mice were treated with KCl alone or KCl (40mM) with 10uM Desipramine, GBR12909 (GBR) or Fluoxetine. Monoamines released into the KH incubation buffer were analyzed by HPLC. *P<0.05, **P<0.01, ****P<0.0001 using Two-Way ANOVA, compared to KCl-treated controls.

**Figure 4.**
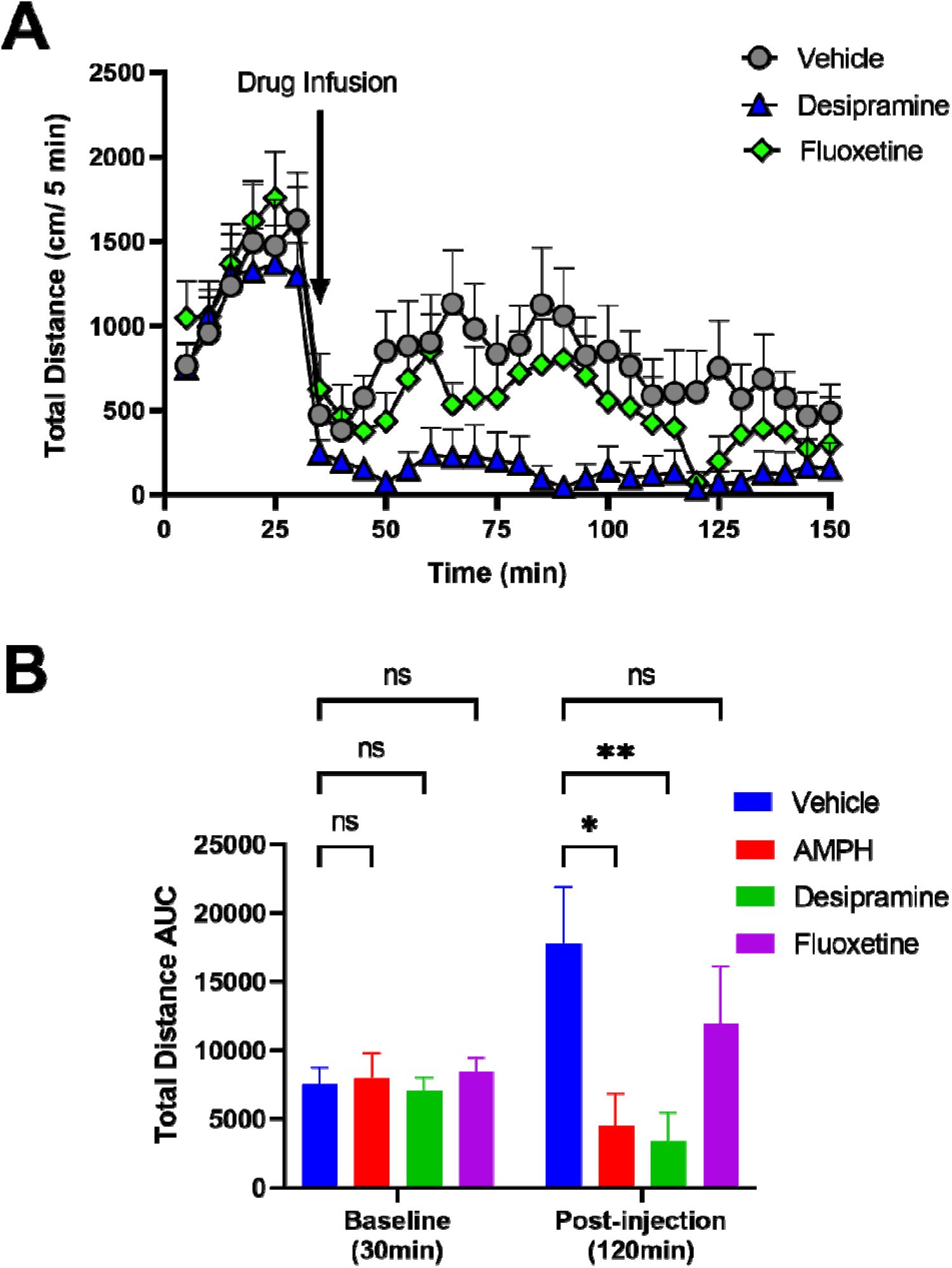
Effect of PFC infusion of monoamine transporter drugs on DAT-KO hyperactivity. Hyperactiv DAT-KO mice were **A**. infused (drug infusion) with vehicle (aCSF) or 4ug/0.5ul of desipramine or fluoxetine, after a 30min baseline, and locomotor activity (cm/5min) was recorded for 120 min post-infusion. **B**. Baseline (cm/30min) and post-infusion (cm/120min) Total distance traveled upon local infusion of desipramine, amphetamine (AMPH) or fluoxetine. n=6-8 mice per group. Pairwise comparisons using Two-way ANOVA, *p<0.05, **p<0.01, ***p<0.001.

Since elevated NE and DA levels in the PFC of DAT-KO mice using NET inhibitors reduced hyperactivity, we next wanted to test the effect of NE depletion in DAT-KO mice. To test this question, we used DBH inhibitor Nepicastat, which depletes NE levels in the brain (Manvich et al., 2013; Schroeder et al., 2013). However, since DBH metabolizes dopamine to norepinephrine, Nepicastat not only depletes NE levels but also enhances DA levels in the PFC (Devoto et al., 2015). Systemic administration (40 mg/kg, i.p.) (**Figure 5 A, B**) or bilateral PFC infusion (4ug/0.5ul/side) (**Figure 5 C, D**) of Nepicastat reduced hyperactivity in DAT-KO mice. HPLC analysis of DAT-KO mice systemically injected with Nepicastat show reduced tissue NE levels and elevated tissue DA levels without altering 5-HT levels in the PFC (**Figure 5C**).

**Figure 5.**
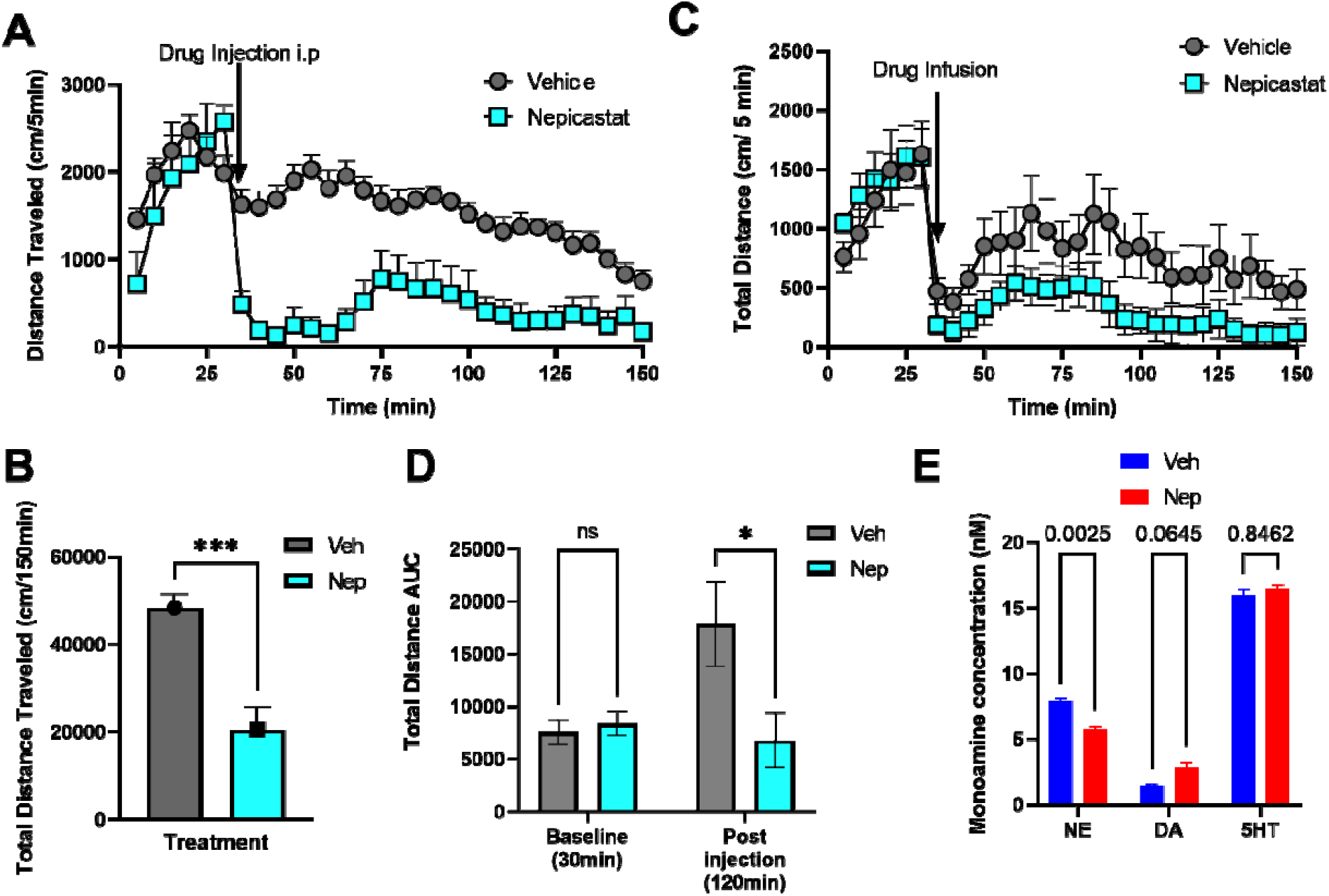
Effect of systemic or local PFC Nepicstat on DAT-KO hyperactivity. Hyperactive DAT-KO mice were systemically injected with **A**. Vehicle or Nepicastat after a 30min baseline, and locomotor activity (cm/5min) recorded for 120min post-injection, and **B**. total distance traveled (cm/150min) calculated. N=6-8 mice per group. Pairwise comparisons using Two-way ANOVA, ***p<0.001. DATKO mice were infused with vehicle or 4ug/0.5ul of Nepicastat, and **C**. locomotor activity (cm/5min) recorded for 120 min or **D**. Total distance traveled baseline (cm/30min) or post-infusion (cm/120min). Pairwise comparisons using Two-way ANOVA, *p<0.05. **E**. DAT-KO PFC tissue was analyzed upon Nepicastat i.p injection, for norepinephrine (NE) or dopamine (DA) or serotonin (5-HT) levels by HPLC. P-values calculated using Two-way ANOVA. Nep compared to vehicle (veh) treated, n=4.

## DISCUSSION

In this study we apply an integrated approach using *ex vivo* efflux techniques, monoamine transporter genetic knockouts, and behavior to understand the molecular and anatomical mechanisms that regulate the phenomenon of paradoxical calming effects of psychostimulants in ADHD therapy. We find that NET inhibitor desipramine but not SERT inhibitor fluoxetine elevates NE and DA levels from PFC tissue slices. Importantly, local PFC infusion of NET blocker desipramine or DBH inhibitor Nepicastat inhibit hyperactivity in DAT-KO mice. These data suggest the elevated PFC DA is a common mechanism that drives the paradoxical calming effect of psychostimulants.

We observe that systemic administration of desipramine and fluoxetine reduces hyperactivity in DAT-KO mice. Although our observations with fluoxetine are consistent with previous findings, they are in contrast to previous findings that the NET inhibitor Nisoxetine did not have any effects in DAT-KO mice (Gainetdinov et al., 1999). However, the dose used in the previous study was 10 mg/kg and other studies have shown that higher doses of Nisoxetine do have behavioral effects in DAT-KO mice (Yamashita et al., 2006), suggesting a role for NET as well in the paradoxical calming effects in DAT-KO mice. These findings are consistent with a role for NET in regulating locomotor activity (Xu et al., 2000). Thus, both NET and SERT might play a role in the paradoxical calming phenomenon of psychostimulants. However, the exact anatomical mechanisms of this phenomenon are not clear. Some studies have suggested that the site of action of psychostimulants is in the PFC and occurs at extremely low doses. These low doses are thought to preferentially act on PFC and not striatal transporters (Berridge et al., 2006). Consistent with these studies we observe that NE and DA levels are elevated in DAT-KO mice in the presence of NET inhibitor desipramine. This phenomenon is mediated by NET since the elevated NE and DA levels are absent in NET-KO mice. However, fluoxetine elevates 5-HT levels and not NE or DA. This is in contrast to previous microdialysis studies showing that fluoxetine increases PFC NE and DA (Bymaster et al., 2002; Tanda et al., 1994). This discrepancy can be explained by the fact that in our study we use tissue slices, where the long-range connectivity of the PFC afferents from midbrain or hindbrain regions are lost, but are maintained in microdialysis studies. However, in our PFC infusion experiments desipramine but not fluoxetine reduces DAT-KO hyperactivity, suggesting that the site of action for fluoxetine is presumably outside the PFC. Recent anatomical mapping studies have suggested that serotonergic raphe nuclei send dense projections to VTA dopamine neurons or locus coeruleus (LC) norepinephrine neurons that project to the medial PFC (Beier et al., 2015; Schwarz et al., 2015; Watabe-Uchida et al., 2012). Thus, it is likely that fluoxetine acts on mesocortical dopamine or LC neurons to elevate PFC DA and NE levels. Overall, two distinct possible mechanisms i.e desipramine in the PFC and fluoxetine in the mid/hind-brain, might regulate DA and NE levels in the PFC. However, DBH inhibitor Nepicastat, which depletes NE but at the same time elevates DA, when locally infused in the PFC also inhibits hyperactivity in DAT-KO mice.

Together these data suggest PFC DA and not NE or 5-HT as a convergent signal for two distinct mechanisms of NET or SERT regulation in the hyperactivity in DAT-KO mice. This notion is consistent with previous data suggesting an inverse role of cortical dopamine and striatal dopamine and locomotor activity (Ventura et al., 2004a; Ventura et al., 2004b). Thus, our data provide a mechanistic insight into the possible mechanisms of paradoxical calming effects of psychostimulants and highlight a predominant role for cortical dopamine in this phenomenon. Our studies have recently identified a possible cortico-striatal-midbrain circuit mechanism for cortical dopamine control of striatal dopamine function and behaviors (Green et al., 2020). Future studies elucidating these circuit mechanisms regulated by this cortical dopamine-dependent phenomenon will provide a possible mechanistic platform for studying regulation of striatal dopamine dysfunction which is a common pathological feature for multiple neurological and psychiatric disorders.

## METHODS

### Animals and Drugs

All mouse studies were conducted in accordance with the NIH guidelines for animal care and use and with an approved animal protocol from the University of Florida Animal Care and Use Committee. The DAT-KO (Giros et al., 1996) and NET-KO (Xu et al., 2000) mice were obtained from Dr. Marc Caron (Duke University). The knockout mice and littermate controls were backcrossed on to a C57BL6/J background for at least 10 generations and are maintained on this background. Amphetamine (Amph, Sigma), desipramine (Sigma), fluoxetine (Sigma), and GBR12909 (R&D systems) were dissolved in saline (behavior) or efflux buffer (efflux assays). Nepicastat (Medkoo, Morrisville, NC) was dissolved in 1% DMSO and 10% hydroxypropyl-cyclodextrin (Sigma), and brought up to volume with saline. Appropriate solvent solutions were used as vehicle controls. All drugs were injected at a volume of 10 ml/kg animal weight.

### Locomotor activity

Locomotor activity was measured in an Accuscan activity monitor (Accuscan Instruments, Columbus, OH) as described previously (Urs et al., 2011). Briefly, locomotor activity was measured at 5 min intervals, and data were analyzed for the total distance traveled in 5 min increments for a total of 150 min or as mentioned in figures. Mice were acclimatized to the activity monitor for 30 min before any drug treatments and drugs were administered after the 30 min acclimatization period, and locomotor activity recorded for an additional 120 min.

### PFC infusions

DAT-KO mice underwent surgery to implant bilateral cannula (Plastics One, 33-gauge, 1 mm) into the PFC using stereotaxic coordinates from a mouse brain atlas (AP: +2.4 mm, ML: 0.5 mm, DV: -1.0 mm from bregma). After at least 1-2 weeks of recovery, the mice were placed in an open-field testing chamber and baseline locomotor activity was assessed for 30 min. Locomotor activity was paused while the mice received a bilateral infusion (0.5 ul/side) of either desipramine (4ug/0.5ul), nepicastat (4ug/0.5ul), amphetamine (100 uM/0.5ul) or vehicle (aCSF or aCSF with 10% cyclodextrin, 1% DMSO) over a 2 min period. Locomotor activity recording was resumed for an additional 120 min.

### *Ex Vivo* Monoamine Release Assay

Wild-type (WT), DAT-KO and NET-KO mice were sacrificed and the brains were quickly removed and immersed in ice-cold Krebs-Henseleit (KH) buffer in the presence of 400 uM ascorbic acid. The composition of KH buffer in mM was: NaCl 116; KCl 3; MgSO4 1; KH2PO4 1.2; NaHCO3 25; D-glucose 11; pH 7.4 saturated with O2/CO2 (95.5% v:v). The brains were sliced into coronal sections (250 um thick) using a vibratome (Leica). Coronal sections of the PFC were collected and kept in ice-cold, oxygenated KH buffer until used for testing. Sections from 4 animals/experiment were equilibrated at 37 C in KH buffer in the presence of 1.8 mM CaCl2, 10 uM pargyline, a MAO inhibitor, and 400 uM ascorbic acid (efflux buffer) for 30 min. After 30 min, the slice were transferred to chambers containing efflux buffer + drug treatment (40 mM KCl +/- 10 uM desipramine, 10 uM GBR 12909, 10 uM Fluoxetine) for 20 min at 37 C. Following incubation with efflux buffer alone or efflux buffer + drug treatment, 0.1 M perchloric acid was added to the supernatants collected from the samples. The supernatants were centrifuged for 5 min at 5000 g.

Monoamine levels were measured using an Eicom HPLC system with an electrochemical detector. Monoamine release was calculated as the amount of catecholamines in the eluate normalized to total protein levels in the tissue samples, and normalized to the KCl-induced control samples.

### Monoamine Tissue Level-HPLC analysis

WT, DAT-KO and NET-KO mice were sacrificed and the brain were placed in a mouse brain matrix. The mouse brain matrix was used to take 1mm coronal sections and punches of the PFC, and dorsal striatum (method described by Salvatore et al., 2012). The punches of each of the brain regions of interest were place on dry ice and stored at -80 C until processed. For processing, tissue samples were placed in ice-cold 0.1 M HClO4 solution and sonicated at 30% of full power. The volume of 0.1 M HClO4 solution was 20 times the weight of PFC and 40 times the weight of dorsal striatum tissue samples. The samples were subsequently centrifuged at 16,000 g for 15 min at 4 C and the supernatants were placed in a separate tube from the protein precipitates. The supernatants were placed in HPLC tubes and monoamine levels were determined by HPLC analysis with electrochemical detection (EICOM).

## ACKNOWLEDGEMENTS

This work was supported by the a NARSAD Young Investigator grant (to NU). We also thank Dr. Marc Caron for providing us with knockout mice.

---

**Figure S1.**
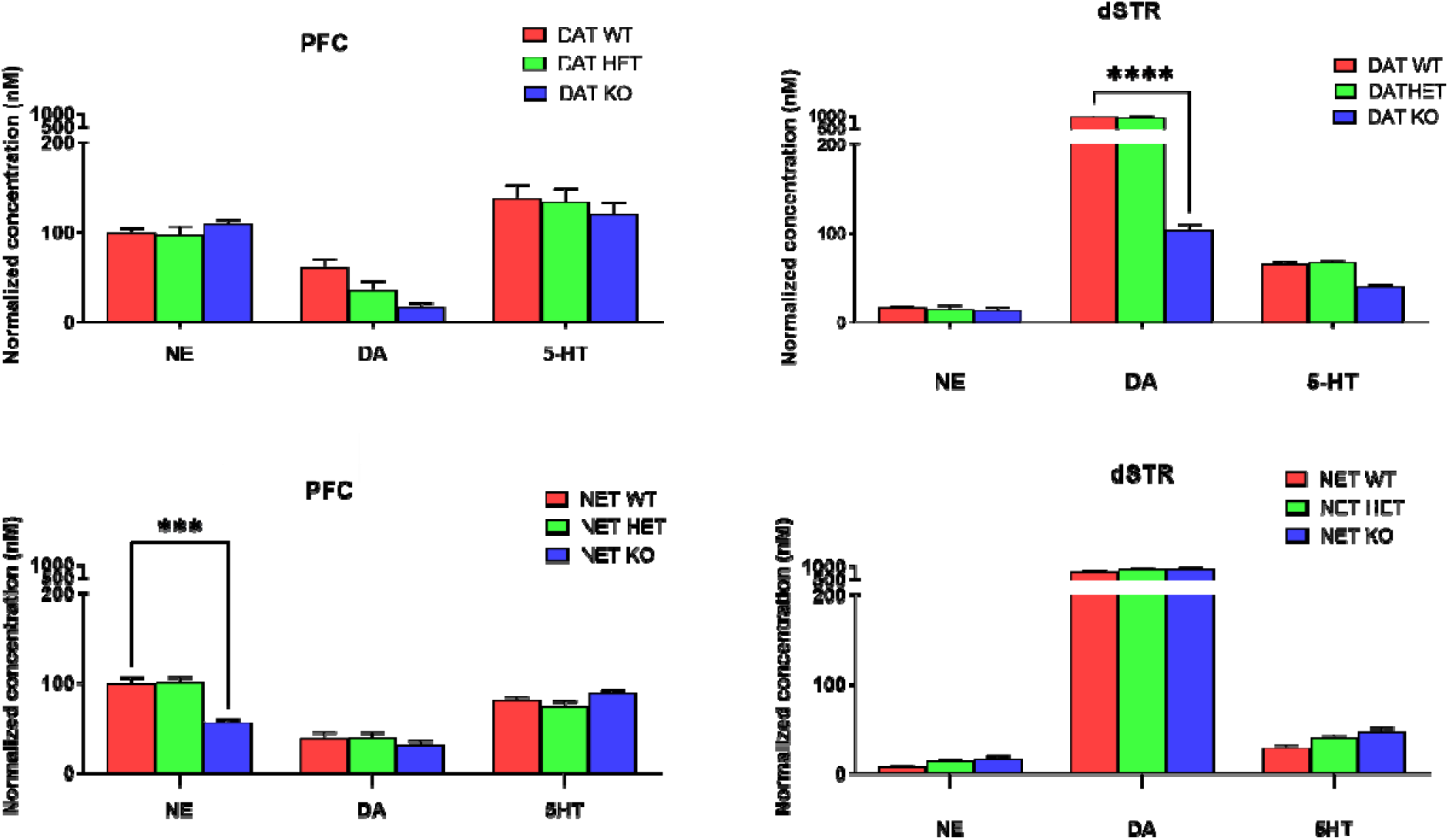
Monoamine tissue levels in DAT-KO and NET-KO mice. DAT-KO or NET KO PFC and dorsal striatum (dSTR) tissue were dissected and analyzed for norepinephrine (NE) or dopamine (DA) or serotonin (5-HT) levels by HPLC. ***P<0.001, ****p<0.0001 using Two-way ANOVA, compared to WT tissue levels n=4.

